# Colour-analyzer: A new dual colour model-based imaging tool to quantify plant disease

**DOI:** 10.1101/2023.08.30.555631

**Authors:** Mackenzie Eli William Loranger, Winfield Yim, Vittorio Accomazzi, Nadia Morales Lizcano, Wolfgang Moeder, Keiko Yoshioka

## Abstract

Despite major efforts over the last decades, the rising demands of the growing global population makes it of paramount importance to increase crop yields and reduce losses caused by plant pathogens. One way to tackle this is to screen novel resistant genotypes and immunity-inducing agents, which must be conducted in a high-throughput manner. Colour-analyzer is a free web-based tool that can be used to rapidly measure the formation of lesions on leaves. Pixel colour values are often used to distinguish infected from healthy tissues. Some programs employ colour models, such as RGB, HSV or L*a*b*. Colour-analyzer uses two colour models, utilizing both HSV (*Hue, Saturation, Value*) and L*a*b* values. We found that the *a* b\** values of the L*a*b* colour model provided the clearest distinction between infected and healthy tissue, while the *H* and *S* channels were best to distinguish the leaf area from the background. By combining the a* and *b\** channels to determine the lesion area, while using the *H* and *S* channels to determine the leaf area, Colour-analyzer provides highly accurate information on the size of the lesion as well as the percentage of infected tissue in a high throughput manner and can accelerate the plant immunity research field.

## Introduction

Over the past 50 years, the agricultural yield of most major crops has matched the rising demand of a growing worldwide population (Simkin et al., 2019). However, recent statistics have reported a troubling plateau in total agricultural output, growing by no more than 1.6% annually (Ray et al., 2013). Current estimates suggest that by the year 2050, the global population will require between a 70% to 100% increase in crop production to meet the demands of a rapidly expanding population, reflecting social shifts towards plant-based products (Tilman & Clark, 2015) and to keep up with the increase in living standards around the globe.

Plant disease still represents one of the major causes of crop loss in modern agriculture. Narrow genetic diversity of large-scale monocultures paired with close growth cycles enables the proliferation of disease leading to a near 30% loss in annual crop yield (Otani et al., 2019; Savary et al., 2019). The fungal necrotrophic pathogen, *Botrytis cinerea,* commonly known as gray mold, is a major concern for agriculture, causing widespread crop loss in a broad range of species and therefore has a destructive economic impact (Fillinger & Elad, 2015, Sivakumar et al., 2016). Worldwide losses due to *B. cinerea* reach over $10 billion annually (Weiberg et al., 2013). *B. cinerea* causes necrosis through the secretion of toxins and enzymes to ultimately feed on dead host cells (van Kan et al., 2014). Currently, chemical control through fungicides is the principal method of reducing *B. cinerea* diseases (Elad et al., 2007). However, when under selective pressure from fungicides, *B. cinerea* can rapidly develop resistance, therefore leading to failure of disease control (Saito et al., 2019). Furthermore, the possible negative impacts of fungicides on the environment and human health are of serious concern (Droby et al., 2009). Therefore, it is necessary to understand *B. cinerea* pathogenesis and form new strategies to control its effect on plants.

Research into plant-pathogen interactions provide an important avenue towards developing strategies to mitigate pathogen-related crop losses. Regardless of the approach taken to make crops more pathogen-resilient, either through the application of a novel compound or the engineering of a genetically altered plant species, an efficient and high throughput assay is instrumental to measure the outcome of an infection. Quantitative measurement of disease and resistance have been well established, including the measure of bacterial load (Pal et al., 2021), the measurement of reactive oxygen species (ROS) production (Yu et al., 2017), the quantification of marker genes, and the measurement of foliar disease symptoms. The measurement of foliar disease symptoms is one of the simplest forms of disease quantification, as it requires minimal training and can be relatively high throughput when compared to other methods. Depending on the pathogen, the development of foliar symptoms can range from the appearance of chlorotic tissue (as seen in the *Pseudomonas syringae* pv. *tomato* DC3000 – *Arabidopsis thaliana* pathosystem) to the development of necrotic lesions (as seen in the interaction between *Magnaporthe oryzae* – *Oryza sativa*).

Image analysis tools exist that can facilitate fast and easy analysis of foliar disease symptoms, such as PIDIQ (Laflamme et al., 2016) for the development of pathogen induced leaf chlorosis. On the other hand, the quantification of fungal lesions (i.e., necrotic tissue) has primarily been done manually using image processing programs like ImageJ (Schneider et al., 2012). ImageJ can be used to simply measure lesion diameters (Li et al., 2020), to set grid areas to be counted (Xu et al., 2023) or even to approximate lesion area based on pixel values using the colour thresholding tool (Kharisma et al., 2022). While the use of image processing programs like ImageJ to quantify disease symptoms has been the standard in the field for many years, it is a relatively time-consuming process, and ill equipped for accurate measurements of irregularly shaped lesions. Other methods such as the use of the hardware YMJ-C smart leaf area meter (Su et al., 2023) and the software Adobe photoshop (Fu et al., 2022) have also been used, but can be cost prohibitive. To this end we sought to develop a tool to rapidly quantify fungal lesions, and for this study we focused on those caused by *Botrytis cinerea* on tomato leaves.

Colour-analyzer is a free web-based tool that can be used to rapidly measure the formation of singular lesions on leaves. By measuring the area of the lesion along with the area of the leaf, Colour-analyzer is not only able to provide information on the size of the lesion, but also the percentage of infected tissue. This provides a quantitative assessment of disease symptoms in a high-throughput manner, allowing for the rapid quantification of disease development while maintaining high precision and accuracy.

## Results and discussion

### Colour models

Whether processing lesion quantification through manual or semi-automated methods, pixel colour values are often used to distinguish infected tissues from healthy tissues. These programs employ colour models, such as RGB, HSV or L*a*b* to assign a set of numerical values to each pixel of an image. The user can then define the threshold within these models to delineate what constitutes healthy versus infected tissues, which can be computed into a total number of pixels and then finally be converted to a relative area or a percentage of infected tissue. The choice of colour model can be imperative when trying to properly discern healthy from infected tissues, especially in a relatively high throughput fashion. The RGB (*Red, Green, Blue*) colour model is usually used for the display optimization of screens and lacks information regarding the illumination of an image (Zajc et al., 2003). Unlike RGB, HSV (*Hue, Saturation, Value*) uses the *Value* portion to help discern changes in colour that occur from shadows or uneven illumination, thus providing a much more reliable way to process images when replicable illumination cannot be guaranteed. When using the RGB model, the separate channel values across a single colour panel with uneven illumination would be different, whereases when using the HSV model, the *Hue* component would be very similar for the whole panel, with the impact of the illumination primarily influencing the *Value* portion of the output. This is an important consideration when designing a software for user accessibility and versatility across a diverse set of plant pathosystems. However, the HSV model is limited in its *Hue* value, as it is a circular value plotted numerically from 0° to 360°, and thus for statistical purposes and computation, circular statistics must be used (Ibraheem et al., 2012). In addition, the *Hue* is defined in 60° slices, in which the relationship between lightness, *Value* and chroma to *R, G, B* depends on each unique slice. This definition introduces discontinuities when we compare slices of the HSV model and can thus make it challenging to use when trying to set thresholds.

The L*a*b* colour model uses *L\** to define the lightness of the object, *a\** for the ratio of red to green, while *b\** defines the ratio of blue to yellow. When plotted, these values provide a 3-dimensional overview, where the *L\** value acts as the Z axis while the *a\** and *b\** values form the X and Y axes respectively, with the sign (negative or positive) of the value inferring directionality towards the opposing colour. Because of this, the L*a*b* colour model is considered a linear model and becomes far superior in terms of computation and statistics compared to the circular nature of HSV. Using these two colour models that include an illumination value (HSV and L*a*b*) we tested their ability to discern the *Botrytis cinerea* lesions on the leaves of tomato plants (Fig. 1).

**Fig 1.**
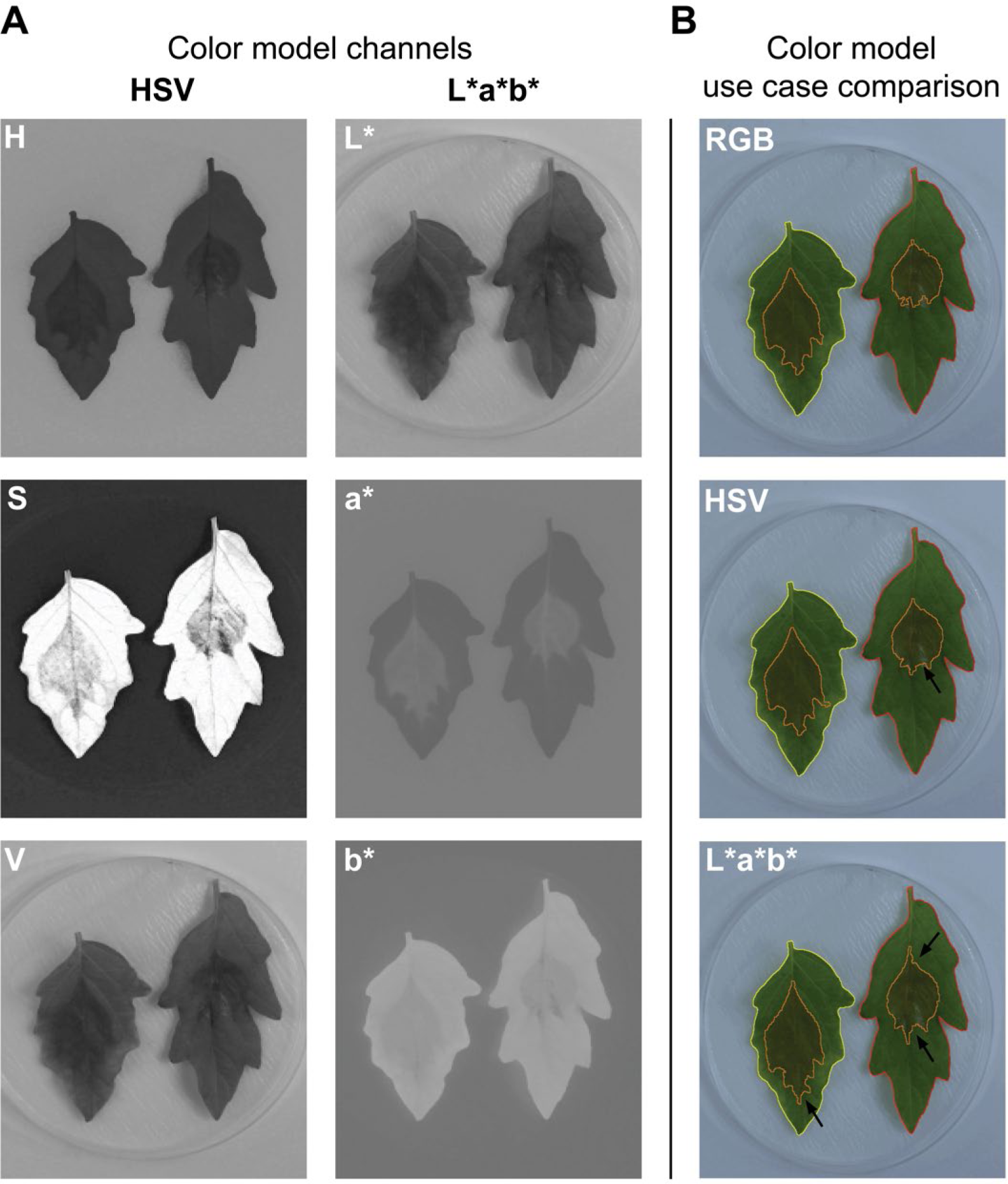
**A.** Separation of the color model channels that make up the HSV and L*a*b* colour models. **B.** Use of the three color models to identify the area of lesion using the same image as input. The black arrows indicate the areas that have become more accurate when compared to the use of the previous model.

Using the same image inputs, it was determined that using the *a* b\** values of the L*a*b* colour model provided the clearest distinction between infected and healthy tissue. This was compared against the use of the *H* and *S* values of the HSV colour model and the combination of all three RGB channels (Fig. 1B). However, it was also noted the *H* and *S* channels where sufficient in determining the leaf area from the background, most notably the saturation channel which shows a strong contrast between the leaf and the uniform background that surrounds it. Therefore Colour-analyzer was built to use the a* and *b\** channels for determining the lesion area, while using the *H* and *S* channels to determine the leaf area.

### Mathematical morphology

One of the prominent issues when using color thresholding to isolate infected tissues is the unwanted selection of healthy tissue and background pixels. This exaggerates the perceived lesion area and may lead to un-reliable quantification of infected tissue, which may especially be an issue when comparing across samples that may have minute variations in basal leaf coloration (e.g., when dealing with mutants that may have altered basal leaf color compared to wild type plants). To overcome this issue, Colour-analyzer uses morphological processing to remove as much background noise as possible. The program leverages a hole-filling algorithm, which is built on the assumption that the hole is the background region surrounded by an un-interrupted border of foreground elements (Ledda et al., 2006). This means that the program will be forced to determine a lesion area that is a single complete region, rather than allowing the thresholding to isolate unwanted pixels (Fig. 1). This assumption works extremely well for isolating experimentally produced necrotic lesion areas, as they form as a single continuous space on the leaf, unlike bacterial specking or chlorosis, that may form in a disrupted pattern.

The use of the Colour-analyzer program dramatically increases the precision of lesion measurement and can reduce overestimations by an average of 20% (Fig. 2). When comparing the use of the L*a*b* model with and without the mathematical morphology, it is also noted that the smaller lesions are more easily overestimated, largely due to the proportional impact of additional pixels to a smaller value. The main pixel area that are misidentified when the mathematical morphology is not employed are regions of shadow surrounding the leaf along with the midvein of the whole leaf, which is often a similar colouration to outer areas of the necrotic tissue. Colour-analyzer completely reduce this noise and provide a consistently more accurate quantification of lesion area.

**Fig. 2.**
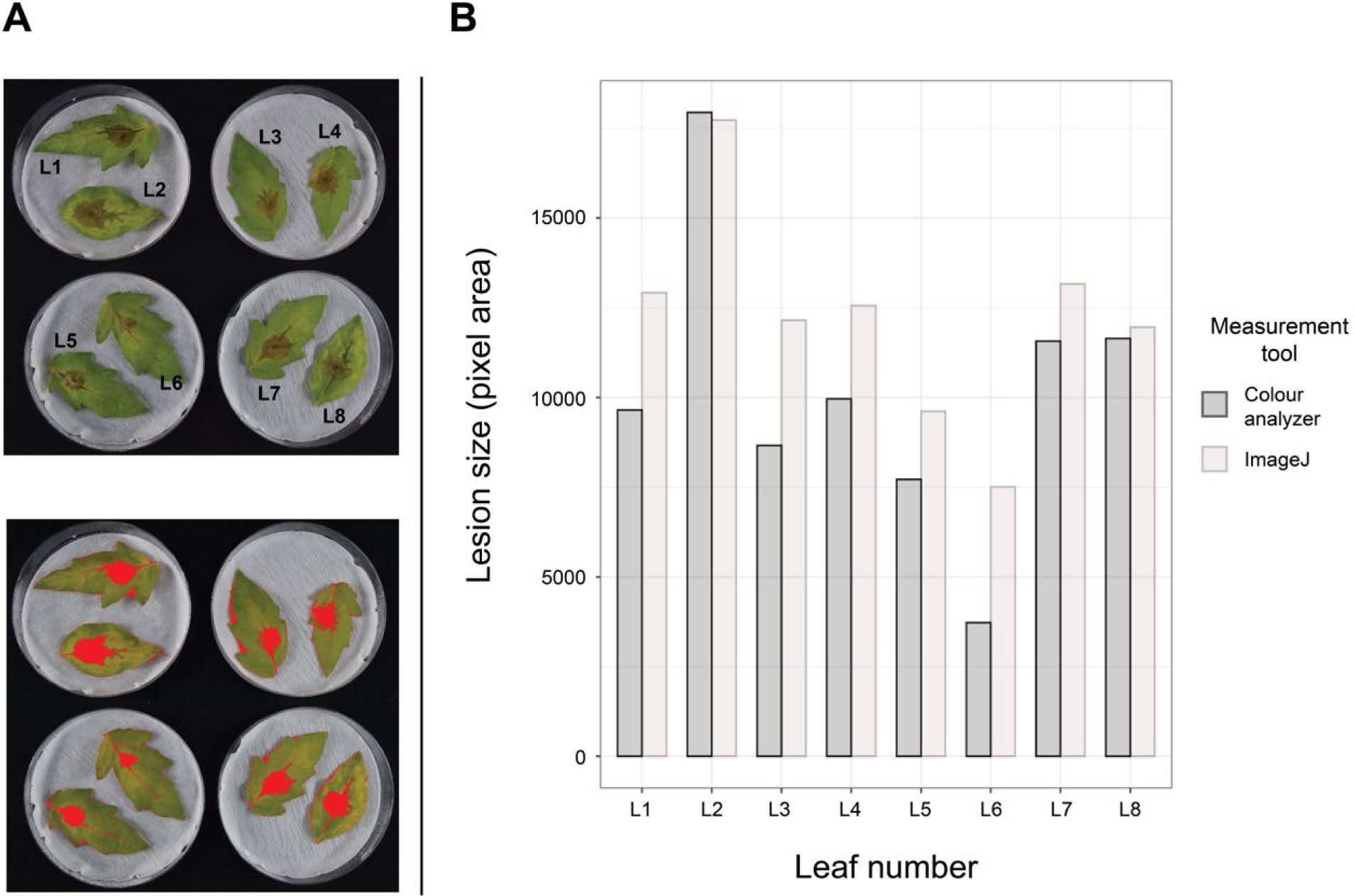
**A.** Lesion identification (red colouration) using the colour thresholding tool in ImageJ; *L\** 0-255, *a\** 119-139, and *b\** 140-173. **B.** Comparison of lesion pixel area when comparing the use of Colour-analyzer versus ImageJ. Leaf number is indicated in the 1^st^ panel of A.

### Colour-analyzer workflow

Colour-analyzer (https://vittorioaccomazzi.github.io/LeafSize/) is a free web-based tool that quantifies the area of necrotic tissue following a fungal infection (Fig. 3). Currently, it is only supported through the free web browser Google Chrome, as it leverages several experimental features unique to Chrome (File system access API, offscreen canvas, PWA and background workers). This tool was developed for automating the measurement of lesions that develop following a detached leaf pathogen assay between *Botrytis cinerea* and *Solanum lycopersicum*. Therefore, some adjustment may be required to use this software for alternate pathosystems.

**Fig. 3.**
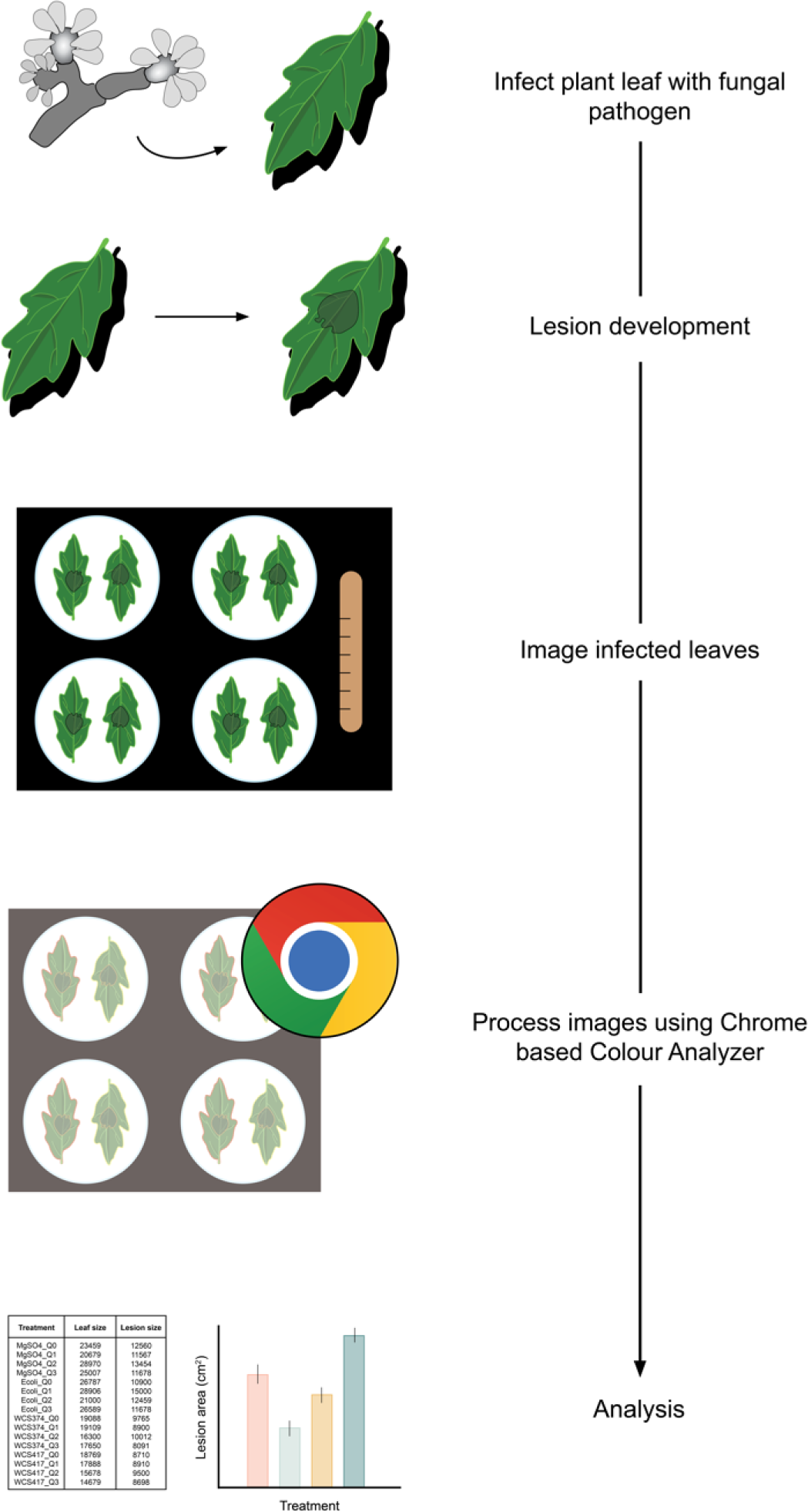
Colour-analyzer workflow outline. Leaves are infected by conidial suspension and left for several days to develop symptoms. Leaves are then batch photographed, in which up to 8 leaves can be photographed together. The web-based program can be run using the Chrome browser and will analyze each leaf individually, producing a CSV file in which each leaf will have both the lesion area and total area indicated. Statistic and graphing can then be done by the user.

Following the infection assay, up to 8 leaves can be imaged at once, in which 2 leaves are placed in each quadrant of the image frame. Either a white or a black background can be used. We recommend using one treatment per image to ease the data sorting following lesion quantification. Photos are taken using a standard digital camera under white light, ideally with even illumination across all leaves. The application allows the selection of an upload folder, from which all captured images should be stored and processed individually. The user interface will prompt the user to select an example leaf region and example lesion region, using the mouse to paint over the region. This will allow the program to gauge what *a\** and *b\** value range is considered infected. The leaf area will be outlined to allow the user to get an output indicating the total leaf area in pixels, which can be used to determine the ratio of infected to uninfected tissue. For this function, the interface has a slider bar which corresponds to the HSV *Hue* and *Saturation* to determine the leaf area. Increasing the hue slider includes more elements into the selection, while increasing the saturation slider will exclude elements from the selection. After clicking the “preview” button, the program will output an outline of the infected tissue and the leaf, the user can adjust the parameters based on the program’s interpretation. Once content with the selection, clicking “next” will allow the program to process all the images using the cut-offs determined in the original image. In most cases, this should be sufficient to determine the lesion area of all images. The user can manually screen through each image to ensure accuracy and discard those that are incorrectly outlined. Then re-select the lesion area and adjust the sliders to adjust the selection. This can be repeated until all images have been processed correctly. The output is a single CSV file ordered by input image, which has been separated into its quadrants, containing both the leaf and lesion area. This data can then be used to compare lesion development across treatments.

### Testing Colour analyzer in tomato and Arabidopsis

To evaluate severity of disease, conventionally, necrotrophic lesions caused by *B. cinerea* are quantified by either manually measuring lesion diameters or lesion areas. These two methods are the simplest approaches to quantify *B. cinerea* symptoms, but, depending on the host plant, there are challenges. For example, in Arabidopsis, *B. cinerea* lesions develop symmetrical, and the lesion diameters correlates with disease severity. However, in tomato the symptoms are often asymmetrical and acentric; thus, standard methods to measure lesion diameters across the midvein can be inaccurate. An alternative method is to measure the entire area of the lesions, which involves manually applying a grid of squares with known dimensions onto the lesion and counting the squares manually. This method is known as the square-counting method (Kvet and Marshall, 1971) and can accurately capture the disease severity and in principle is superior to measuring the diameter. However, this is extremely labor intensive and time-consuming, and thus is not suitable for a high throughput nor large-scale analysis. In addition, manual measurement of lesion areas or even lesion diameters are always at risk of human error and bias.

In this study, we compared lesion area in Arabidopsis and tomato leaves infected with a drop of *B. cinerea* conidia. We compared untreated control plants with plants treated with two potential immunity-inducing agents. Treatment 1 had no effect against *B. cinerea* while treatment 2 lead to significantly smaller *B. cinerea* lesions in both Arabidopsis and tomato plants (Fig. 4). When the lesions were measured along the midvein of Arabidopsis leaves, only treatment 2 showed significantly smaller lesions compared to the control (Fig 4A). This result was consistent with the result where lesion diameters were measured perpendicular to the midvein (Fig. 4B). Although treatment 1 showed different lesion diameters between midvein and perpendicular measurements, the results remained the same where only treatment 2 showed significantly decreased lesions (Fig4 A and B). Thus, on Arabidopsis leaves, due to the formation of circular *B. cinerea* lesions, even measuring lesion diameters on different axes provided similar results. On the other hand, when measuring the lesion diameters along the midvein for tomato leaves, treatment 2 again showed statistically significant decreased *B. cinerea* lesions (Fig. 4C). However, when the same lesions are measured for their diameters perpendicular to the midvein no significant differences were observed (Fig. 4D). Treatment 2 is no longer significantly reduced, and all lesions seem to have nearly halved in diameter. This illustrates that measuring lesion diameters can be extremely inconsistent if the axis of measurement is not defined properly and can be very susceptible to human error.

**Fig. 4.**
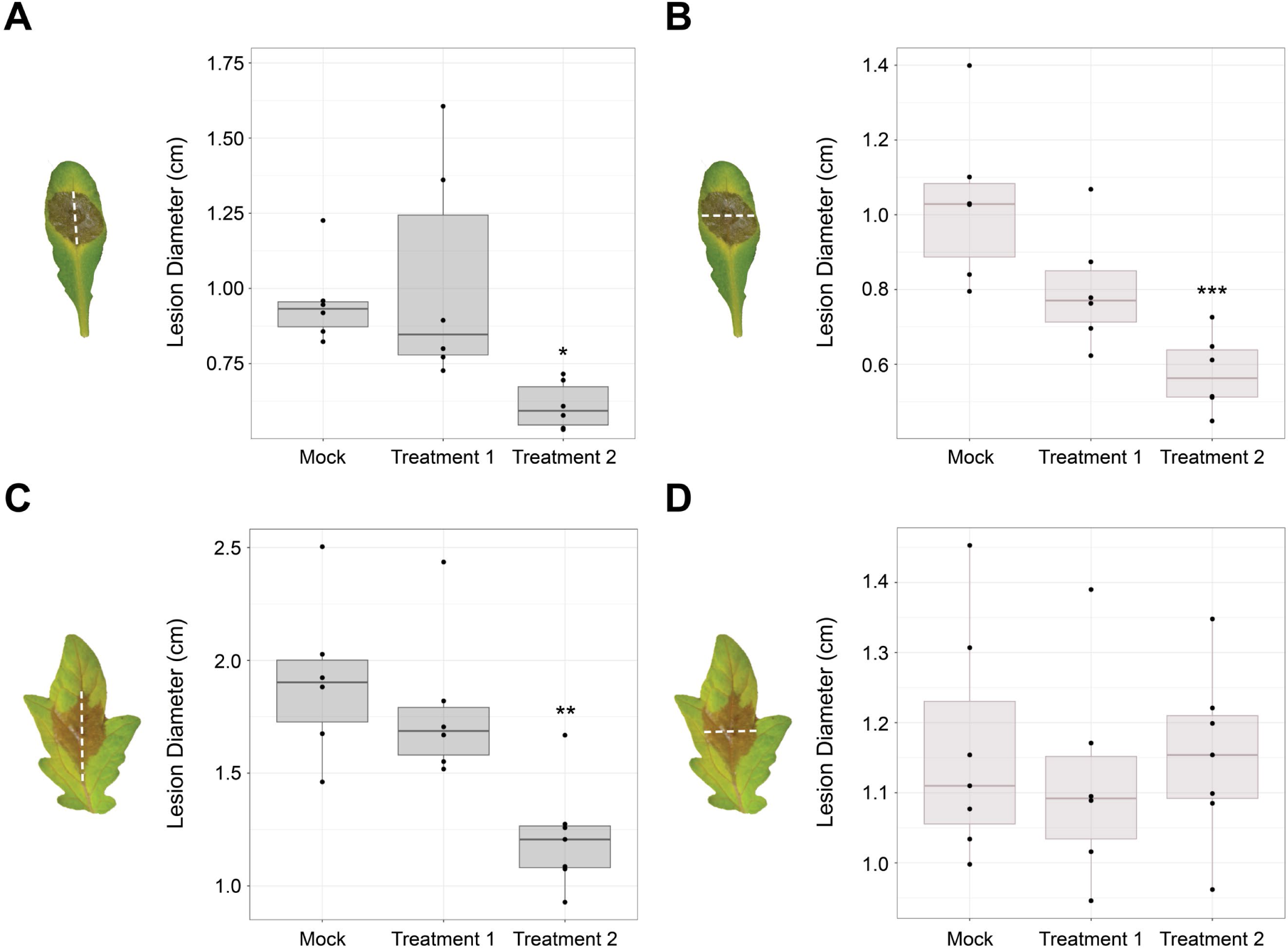
Differences in *B. cinerea* lesion quantification when measuring the diameters horizontally or vertically in Arabidopsis and tomato leaves. Leaves with lesions in this figures are presented to illustrate how lesions were measured in each subgroup. **A.** Measuring lesions on Arabidopsis leaves vertically along the midvein. **B.** Measuring the same subset of leaves as in 2A., but using the horizontal diameter, perpendicular to the midvein. **C.** Measuring lesions on tomato leaves vertically along the midvein. **D.** Measuring the same subset of leaves as in 2C., but using the horizontal diameter, perpendicular to the midvein. Differences between treatment groups and mock were evaluated using a one-way ANOVA followed by a Dunnett’s test post hoc. * ≤ 0.05, ** ≤ 0.01, *** ≤ 0.001. n = 6-8.

### Quantifying immunity

Induced systemic resistance (ISR) is a form of systemic immunity that is conferred through the interactions between ISR-inducing bacteria in the rhizosphere and the plant root. These interactions culminate in primed defence response in the above-ground tissues, largely reliant on the jasmonic acid-associated defence transcription factors, MYC2 and MYB72 (Pozo et al., 2008; Van Der Ent et al., 2008). When testing a bacterial strain’s ability to induce ISR, a pathogen assay is used to measure the level of disease protection, using lesion area as a proxy for the immune response. The gram-negative bacterium *Pseudomonas defensor* WCS374r is a known ISR inducing strain that reduces disease in a variety of plant species, such as radish, Eucalyptus, and rice (De Vleesschauwer et al., 2008; Leemann et al., 1995; Berendsen et al., 2015; Lee-Díaz et al., 2021) but until now, has yet to be experimentally shown to be an elicitor of ISR in *Solanum lycopersicum* (Berendsen et al., 2015). We pre-inoculated young tomato seedlings with either *Pseudomonas defensor* WCS374r, *E. coli* or 10mM MgSO_4_ control). *E. coli* was used a control treatment to mimic the addition of a high titer of bacteria to the rhizosphere. Plants were left to grow in soil for several weeks before leaves were detached, and challenge inoculated. A droplet of *B. cinerea* conidia was added to the midvein of each leaf and the leaves where sealed in petri dishes ensure high relative humidity and promote lesion development. Four-days post infection, images of the infected leaves were captured. Lesions on leaves from plants pre-treated with *Pseudomonas defensor* WCS374r were statistically significantly reduced compared to those from control, while *E. coli* pre-treated plants showed no significant change compared to the mock set (Fig. 5). In this proof-of-concept experiment, we show how this tool can be used to screen treatments that can help plants to defend themselves against disease.

**Fig. 5.**
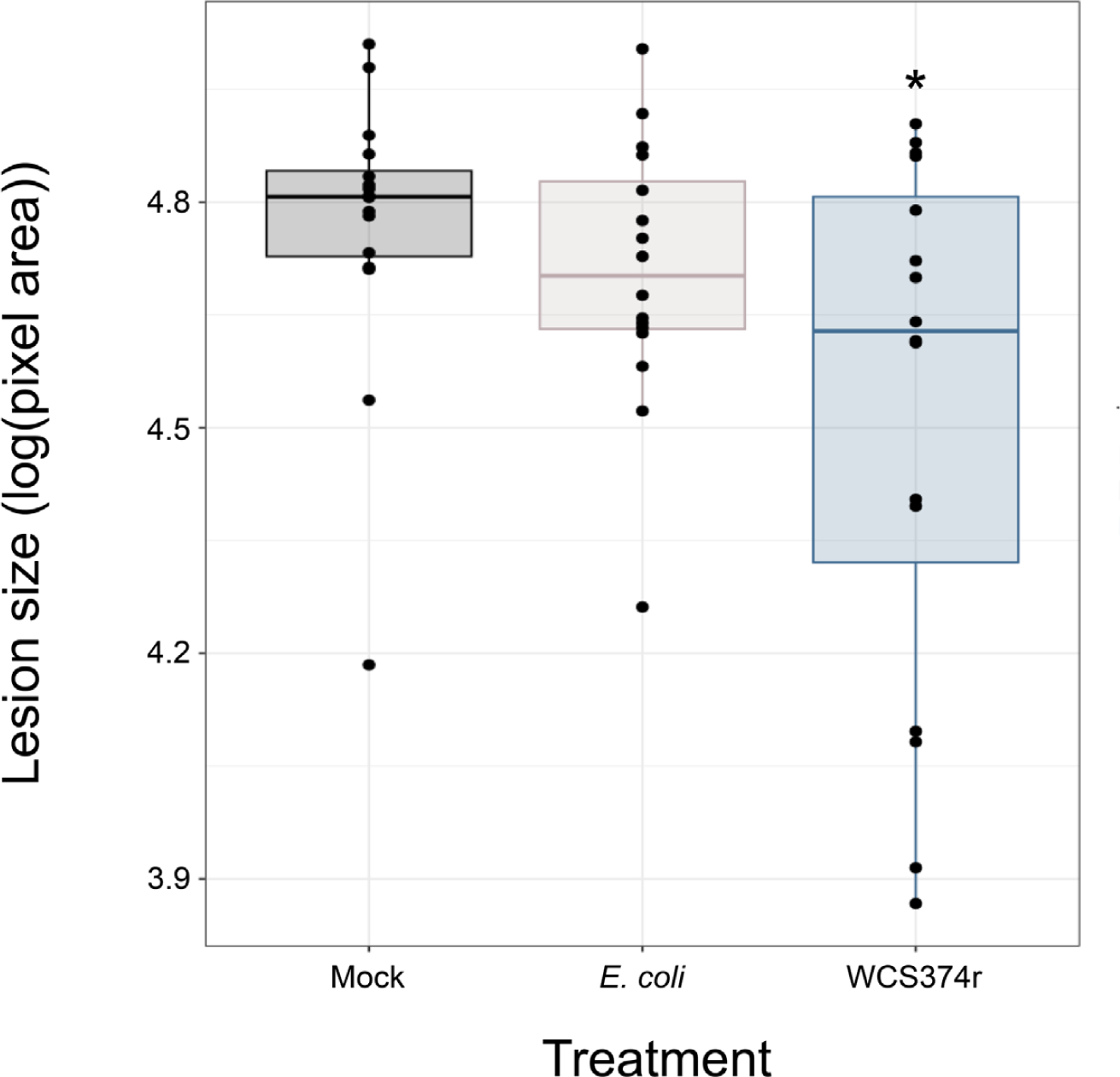
*B. cinerea* lesion area in plants pre-treated with either mock or bacterial solutions. The lesion areas were measured using Colour Analyzer, using pixel area as an output. Differences between treatment groups and mock were evaluated using a one-way ANOVA followed by a Dunnett’s test post hoc. * ≤ 0.05, ** ≤ 0.01. n = 16.

### Conclusion

Colour-analyzer is a free web-based tool for high-throughput screening of lesions on leaves. Here, we present its application for the agriculturally important fungal pathogen, *Botrytis cinerea* in tomato, but this tool can also be utilized for other pathosystems and also for the quantification of discolorations or damage of leaves by abiotic stressors. It can provide very accurate leaf area measurements as well. The application is available at https://github.com/VittorioAccomazzi/LeafSize and is designed to be improved or extended by users. Thus, we developed a new versatile imaging tool that is free for all plant researchers.

## Materials and methods

### Bacterial strains and plant pre-treatment

*Pseudomonas fluorescens* WCS374r and *Escherichia coli* DH5α were grown in Lysogeny broth (LB) at 30°C and 37°C respectively. Bacterial cultures were grown overnight in liquid cultures, before being spun down and resuspended in 10mM MgSO_4_ to an OD_600_ of 0.1. *Solanum lycopersicum* cv. ‘Glamour’ seeds were surface sterilized and germinated on ½ MS 1% sucrose agar plates for 5-days before being transferred to Jiffy-pellets (Jiffy Growing Solutions, Zwijndrecht, the Netherlands) Jiffy-pellets were soaked in 50mL of bacterial inoculum each, for an hour before seed transfer. The plants were grown under 16h of light and 8h of darkness at 22°C. Two weeks post seed sterilization, Jiffy-pellets were transferred to larger pots filled with Sunshine Mix #1 to support the increased nutritional and spatial requirements during the maturation of the plants.

### *Botrytis cinerea* culturing and challenge inoculation

The *Botrytis cinerea* MEE B191 isolate that was used for this experiment was provided by the Canadian Collection of Fungal Cultures (Agriculture and Agri-Food Canada, Ottawa, ON, Canada). The fungus was grown on potato dextrose agar (PDA) for 7 days at room temperature. *B. cinerea* spores (conidia) were collected and vortexed in Sabouraud maltose broth (SMB) before being passed through a Corning cell strainer with a pore size of 100μm, which effectively removes large clumps of conidia and unwanted hyphae from the inoculum. The conidial concentration was adjusted to a final concentration of 2.5 X 10^5^ conidia/mL.

The challenge inoculation was performed by placing a 10μl droplet of the conidia solution in the midvein of detached leaves. A total of 16 leaves from four different plans were used per treatment. The inoculated leaves were houses in sealed petri dishes (2 per dish) lined with water-saturated sterile filter paper to maintain a high humidity environment. The petri dishes were kept in 24h light conditions at room temperature for 3-4 days.

### Image processing and analysis

All images were taken under controlled lighting conditions at a set height, to minimize image distortion and ensure maximal colour consistency. A black cloth was used as the background and a petri dish was added to each corner of the image frame, with a total of 8 leaves being imaged at a time. A ruler was placed within the image frame to allow for the conversion of pixel area to centimeters squared (cm^2^) in the later steps. All images were stored in a single folder and the image files were re-named according to their treatment or condition. The program, available on Github and through the Chrome web browser, semi-automates the process of lesion quantification following the steps below.

Once the image set is loaded onto the program a single image can be used to establish the *a\** and *b\** values used to process the batch by using the cursor to select infected and healthy tissue. Using this approach, the upper and lower bounds of these values can be different between image sets or even between individual images, allowing for a fast and easy way to adjust thresholds between the sets. The sliders at the top are used to help isolate the leaf region, which can also be modified between images. Once the program has finished processing the images, the user can identify images with improper selections, and the selection process can be re-preformed on this subset of images. This process can be repeated as many times as needed. Once all the images are accepted the program will output a CSV file which can be downloaded. The first column is the image name, which will be the file name and either Q0, Q1, Q2 or Q4 depending on the quadrant. The other columns will include the leaf area and the pathogen/lesion area. This data can be easily modified for statistical or graphing purposes.

## Acknowledgments

This work was supported by a NSERC Strategic Partnership Grant to KY, and an Ontario Graduate Scholarship to MEWL.

## Notes

### Competing Interest Statement

The authors have declared no competing interest.

## Literature cited

Berendsen, R. L., van Verk, M. C., Stringlis, I. A., Zamioudis, C., Tommassen, J., Pieterse, C. M. J., & Bakker, P. A. H. M. (2015). Unearthing the genomes of plant-beneficial Pseudomonas model strains WCS358, WCS374 and WCS417. BMC Genomics, 16(1). 10.1186/s12864-015-1632-z

Droby, S., Wisniewski, M., Macarisin, D., & Wilson, C. (2009). Twenty years of postharvest biocontrol research: Is it time for a new paradigm? Postharvest Biology and Technology, 52(2), 137–145. 10.1016/j.postharvbio.2008.11.009

Elad, Y., & Stewart, A. (2007). Microbial control of botrytis spp. In Botrytis: Biology, Pathology and Control. Springer. 223–241. 10.1007/978-1-4020-2626-3_13

Fillinger, S., & Elad, Y. (2015). Botrytis-the Fungus, the Pathogen and its Management in Agricultural Systems. https://www.researchgate.net/publication/287645053

Fu, Y., Li, J., Wu, H., Jiang, S., Zhu, Y., Liu, C., Xu, W. J., Li, Q., & Yang, L. (2022). Analyses of Botrytis cinerea-responsive LrWRKY genes from Lilium regale reveal distinct roles of two LrWRKY transcription factors in mediating responses to B. cinerea. Plant Cell Reports, 41(4), 995–1012. 10.1007/s00299-022-02833-6

Ibraheem, N. A., Hasan, M. M., Khan, R. Z., & Mishra, P. K. (2012). ARPN Journal of Science and Technology:: Understanding Color Models: A Review. ARPN Journal of Science and Technology, 2(3). http://www.ejournalofscience.org

Kharisma, A. D., Arofatullah, N. A., Yamane, K., Tanabata, S., & Sato, T. (2022). Regulation of defense responses via heat shock transcription factors in Cucumis sativus L. against Botrytis cinerea. Journal of General Plant Pathology, 88(1), 17–28. 10.1007/s10327-021-01041-6

Květ, J., Marshall, J.K. (1971). Assessment of leaf area and other assimilating plant surfaces. In: Šesták, Z., Čatský, J., Jarvis, P.G. (ed.): Plant Photosynthetic Production. Manual of Methods. Dr W. Junk Publ., The Hague. 517–555.

Laflamme, B., Middleton, M., Lo, T., Desveaux, D., & Guttman, D. S. (2016). Image-based quantification of plant immunity and disease. Molecular Plant-Microbe Interactions, 29(12), 919–924. 10.1094/MPMI-07-16-0129-TA

Ledda, A., Luong, H. Q., Philips, W., De Witte, V., & Kerre, E. E. (2006). Image Interpolation using Mathematical Morphology.

Lee Díaz, A. S., Macheda, D., Saha, H., Ploll, U., Orine, D., & Biere, A. (2021). Tackling the context-dependency of microbial-induced resistance. In Agronomy (Vol. 11, Issue 7). MDPI AG. 10.3390/agronomy11071293

Li, R., Sheng, J., & Shen, L. (2020). Nitric oxide plays an important role in β-aminobutyric acid-induced resistance to botrytis cinerea in tomato plants. Plant Pathology Journal, 36(2), 121–132. 10.5423/PPJ.OA.11.2019.0274

Otani, S., Challinor, V. L., Kreuzenbeck, N. B., Kildgaard, S., Krath Christensen, S., Larsen, L. L. M., Aanen, D. K., Rasmussen, S. A., Beemelmanns, C., & Poulsen, M. (2019). Disease-free monoculture farming by fungus-growing termites. Scientific Reports 2019 9:1, 9(1), 1–10. 10.1038/s41598-019-45364-z

Pal, G., Mehta, D., Singh, S., Magal, K., Gupta, S., Jha, G., Bajaj, A., & Ramu, V. S. (2021). Foliar Application or Seed Priming of Cholic Acid-Glycine Conjugates can Mitigate/Prevent the Rice Bacterial Leaf Blight Disease via Activating Plant Defense Genes. Frontiers in Plant Science, 12. 10.3389/fpls.2021.746912

Pozo, M. J., Van Der Ent, S., Van Loon, L. C., & Pieterse, C. M. J. (2008). Transcription factor MYC2 is involved in priming for enhanced defense during rhizobacteria-induced systemic resistance in Arabidopsis thaliana. New Phytologist, 180(2), 511–523. 10.1111/j.1469-8137.2008.02578.x

Ray, D. K., Mueller, N. D., West, P. C., & Foley, J. A. (2013). Yield Trends Are Insufficient to Double Global Crop Production by 2050. PLOS ONE, 8(6), e66428. 10.1371/JOURNAL.PONE.0066428

Saito, S., Michailides, T. J., & Xiao, C. L. (2019). Fungicide-resistant phenotypes in Botrytis cinerea populations and their impact on control of gray mold on stored table grapes in California. European Journal of Plant Pathology, 154(2), 203–213. 10.1007/s10658-018-01649-z

Savary, S., Willocquet, L., Pethybridge, S. J., Esker, P., McRoberts, N., & Nelson, A. (2019). The global burden of pathogens and pests on major food crops. Nature Ecology & Evolution 2019 3:3, 3(3), 430–439. 10.1038/s41559-018-0793-y

Schneider, C. A., Rasband, W. S., & Eliceiri, K. W. (2012). NIH Image to ImageJ: 25 years of image analysis. In Nature Methods (Vol. 9, Issue 7, pp. 671–675). 10.1038/nmeth.2089

Simkin, A. J., López-Calcagno, P. E., & Raines, C. A. (2019). Feeding the world: improving photosynthetic efficiency for sustainable crop production. Journal of Experimental Botany, 70(4), 1119–1140. 10.1093/JXB/ERY445

Su, K., Zhao, W., Lin, H., Jiang, C., Zhao, Y., & Guo, Y. (2023). Candidate gene discovery of Botrytis cinerea resistance in grapevine based on QTL mapping and RNA-seq. Frontiers in Plant Science, 14. 10.3389/fpls.2023.1127206

Tilman, D., & Clark, M. (2015). Food, agriculture & the environment: Can we feed the world & save the earth? Daedalus, 144(4), 8–23. 10.1162/DAED_A_00350

Van Der Ent, S., Verhagen, B. W. M., Van Doorn, R., Bakker, D., Verlaan, M. G., Pel, M. J. C., Joosten, R. G., Proveniers, M. C. G., Van Loon, L. C., Ton, J., & Pieterse, C. M. J. (2008). MYB72 is required in early signaling steps of rhizobacteria-induced systemic resistance in arabidopsis. Plant Physiology, 146(3), 1293–1304. 10.1104/PP.107.113829

van Kan, J. A. L., Shaw, M. W., & Grant-Downton, R. T. (2014). Botrytis species: Relentless necrotrophic thugs or endophytes gone rogue? Mol Plant Pathol. 15:957–61. 10.1111/mpp.12148

Weiberg, A., Wang, M., Lin, F.-M., Zhao, H., Zhang, Z., Kaloshian, I., Huang, H.-D., & Jin, H. (2013). Fungal Small RNAs Suppress Plant Immunity by Hijacking Host RNA Interference Pathways. Science, 342, 118–123. https://api.semanticscholar.org/CorpusID:14989348

Xu, J., Yan, D., Chen, Y., Cai, D., Huang, F., Zhu, L., Zhang, X., Luan, S., Xiao, C., & Huang, Q. (2023). Fungicidal activity of novel quinazolin-6-ylcarboxylates and mode of action on Botrytis cinerea. Pest Management Science, 79(9), 3022–3032. 10.1002/ps.7477

Yu, X., Feng, B., He, P., & Shan, L. (2017). From Chaos to Harmony: Responses and Signaling upon Microbial Pattern Recognition. Annual Review of Phytopathology, 55(1), 109–137. 10.1146/annurev-phyto-080516-035649

Zajc, Baldomir., Tkalčič, Marko., Institute of Electrical and Electronics Engineers., & Institute of Electrical and Electronics Engineers. Region 8. (2003). The IEEE Region 8 EUROCON 2003: computer as a tool: proceedings: 22-24 September 2003, Faculty of Electrical Engineering, University of Ljubljana, Ljubljana, Slovenia. IEEE.

